# Application of the fluorescence-activating and absorption-shifting tag (FAST) for flow cytometry in methanogenic archaea

**DOI:** 10.1101/2022.08.04.502898

**Authors:** Norman Adlung, Silvan Scheller

**Affiliations:** School of Chemical Engineering, Department of Bioproducts and Biosystems, Aalto University, 02150 Espoo, Finland; VTT Technical Research Centre of Finland Ltd., 02044 Espoo, Finland

**Keywords:** Methanogenic archaea, Fluorescence, Flow Cytometry, FAST, *Methanosarcina*

## Abstract

Methane-producing archaea play a crucial role in the global carbon cycle and are used for biotechnological fuel production. Methanogenic model organisms such as *Methanococcus maripaludis* and *Methanosarcina acetivorans* are biochemically characterized and can be genetically engineered using a variety of molecular tools. Methanogens’ anaerobic lifestyle and autofluorescence, however, restrict the use of common fluorescent reporter proteins (e.g., GFP and derivatives) which require oxygen for chromophore maturation. Here, we employ the tandem activation and absorption-shifting tag protein 2 (tdFAST2) which is fluorescent when the cell-permeable fluorescent ligand (fluorogen) 4-hydroxy-3,5-dimethoxybenzylidene rhodanine (HBR-3,5DOM) is present. tdFAST2 expression in *M. acetivorans* and *M. maripaludis* is not cytotoxic and tdFAST2:HBR-3,5DOM fluorescence can be clearly distinguished from the autofluorescence. In flow cytometry experiments, mixed methanogen cultures can be clearly distinguished which allows high-throughput investigations of dynamics within single and mixed cultures.

**Importance:** Methane-producing archaea play an essential role in the global carbon cycle and have a high potential for biotechnological applications such as biofuel production, carbon dioxide capture, and in electrochemical systems. The oxygen sensitivity and high autofluorescence hinder the use of common fluorescent proteins to study methanogens. By using the tdFAST2:HBR-3,5DOM fluorescence, which is functional also under anaerobic conditions and distinguishable from the autofluorescence, real-time reporter studies and high-throughput investigation of dynamics within (mixed) cultures via flow cytometry are possible. This will accelerate the exploitation of the methanogens’ biotechnological potential.

## Introduction

Methanogenic archaea are responsible for up to 70% of the methane emitted globally (1). Besides their relevance for the global carbon cycle, methanogens have a high potential for biotechnological applications. Applications include (i) production of biogas and biofuel (2), (ii) treatment of solids and sewage water (3), (iii) carbon dioxide capture (4, 5), and (iv) storage systems for excess electricity via bioelectrochemical systems (6). Genetically tractable methanogens include *Methanococcus maripaludis* and *Methanosarcina* species. Genetic tools for methanogens include plasmid-expression systems, genome modification via homologous recombination and CRISPR/Cas, inducible gene expression, and reporter systems (7–9).

The oxygen sensitivity of methanogens limits the use of common fluorescent proteins. Fluorescent proteins with a barrel-like structure (e.g. GFP and mCherry) require oxygen for chromophore maturation and are not fluorescent under anaerobic conditions (10). The use of fluorescent proteins in methanogens is also hindered by the cells’ autofluorescence (emission maximum at 480 nm), which originates from oxidized coenzyme F420 after excitation at 420 nm (11). Consequently, valuable tools such as protein localization via fluorescence microscopy, high-throughput measurement of protein accumulation via flow cytometry, and fluorescence-activated cell sorting (FACS) are underexplored for methanogens. In recent years, novel tools to replace oxygen-dependent fluorescent proteins under anaerobic conditions were developed. These tools employ different mechanisms, for example (i) flavin mononucleotide-based fluorescent proteins, (ii) fluorescence activating proteins which require reversible binding of a fluorogenic ligand, and (iii) self-labelling proteins that require covalent binding to non-fluorogenic ligands (12, 13). Recently, a fluorescence activating protein was successfully applied in the methanogen *M. maripaludis* (14).

Fluorescence-activating proteins such as FAST (*fluorescence-activating and absorption-shifting tag*) and its improved version FAST2 are small, engineered proteins that become fluorescent upon binding to a fluorogenic ligand (fluorogen) (15, 16). FAST and FAST2 can be N- or C-terminally fused to a protein of interest, expressed as single protein or as a tandem protein to increase fluorescence (tdFAST/tdFAST2). Different fluorogens were developed; they are 4-hydroxybenzylidene rhodanine derivatives and determine the excitation/emission wavelength of the FAST:fluorogen fluorescene upon interaction. FAST can also be used to study *in vivo* protein-protein interaction where the N- and C-terminal parts of FAST are fused to the proteins of interest (splitFAST). FAST:fluorogen fluorescence can be detected when the two proteins of interest interact and bring the N- and C-terminal FAST parts in close proximity (17). Here, we apply the FAST reporter to the methanogenic model organism *M. acetivorans* and we demonstrate that the fluorescent reporter provides reliable results in flow cytometry experiments.

## Results

### tdFAST2:HBR-3,5DOM fluorescence in methanogens

Plasmids for constitutive expression of tdFAST2 in *M. acetivorans* (pNB730::tdFAST2) and *M. maripaludis* (pMEV4::tdFAST2) were generated and transformed into methanogens. Cells transformed with the original (empty) vectors served as control strains. Liquid cultures were analyzed using a microplate reader under aerobic conditions. A washing step was introduced to eliminate background fluorescence from the medium. When the fluorogen HBR-3,5DOM was present, tdFAST2-producing *M. acetivorans* and *M. maripaludis* cells showed a fluorescence peak around 590 nm when excited at 515 nm (Fig. 1a). This fluorescence peak was not detected when the fluorogen or tdFAST2 was absent. Thus, tdFAST2:HBR-3,5DOM fluorescence can be clearly separated from methanogens’ autofluorescence. Further analysis showed that tdFAST2:HBR-3,5DOM fluorescence can be excited upon a wide range of wavelengths ranging from 480 nm to 550 nm (Fig. 1b). This is in contrast to the fluorogen HMBR, which has a rather narrow excitation range around 490 nm (Supplementary Fig. S1). In further experiments, standard settings of 515 nm for excitations and 590 nm for emission were used for detection of tdFAST2:HBR-3,5DOM fluorescence.

**Figure 1.**
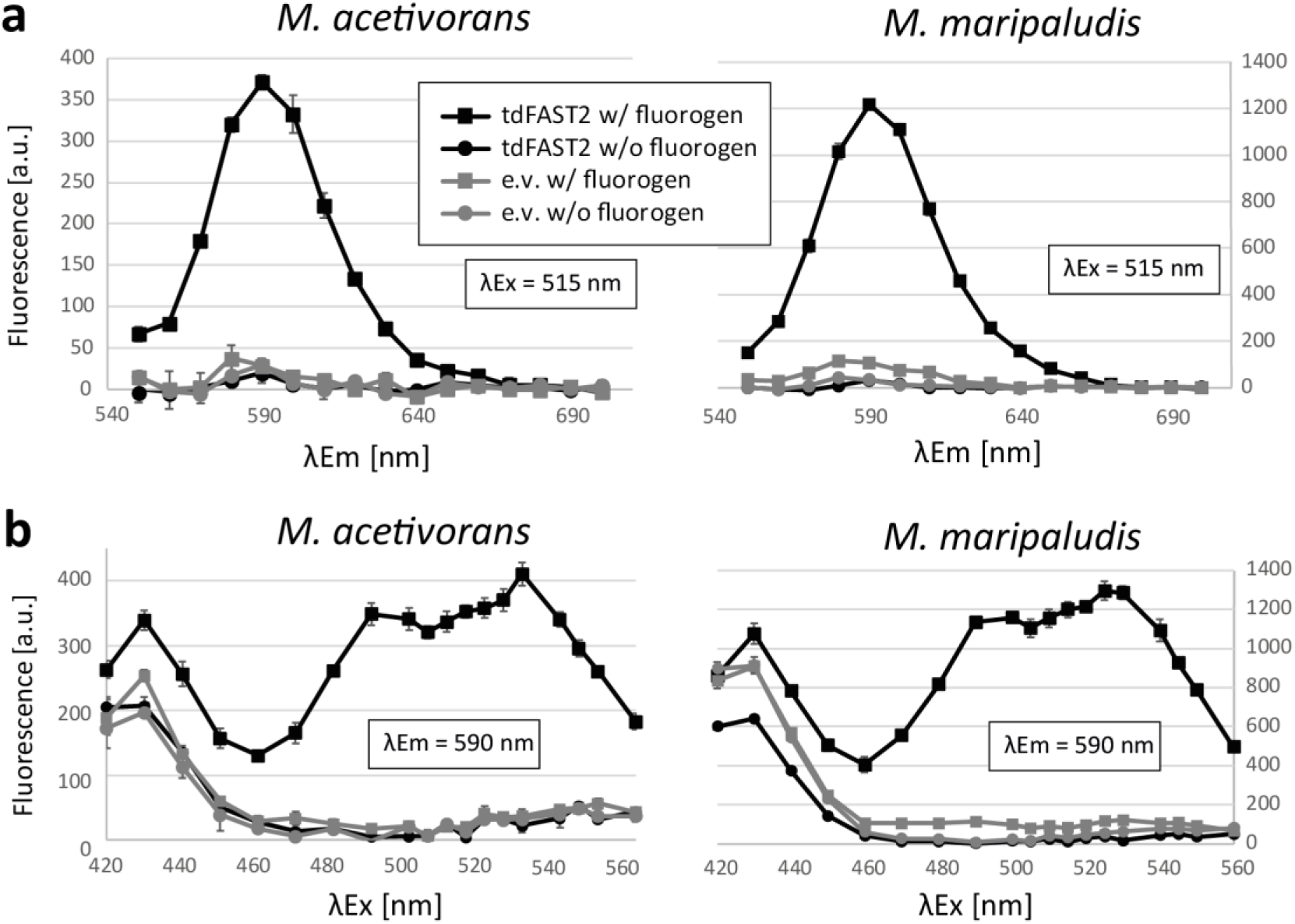
tdFAST2-expressing methanogens show a specific fluorescence in the presence of HBR-3,5DOM. Fluorescence of *M. acetivorans* and *M. maripaludis* cells is shown. **a**) Fluorescence spectrum upon excitation at 515 nm. **b**) Fluorescence at 590 nm when different excitation wavelengths are applied. Cells expressing tdFAST2 (tdFAST2) or control cells carrying an empty vector construct (e.v.) were analysed in presence (w/ fluorogen) or absence (w/o fluorogen) of HBR-3,5DOM. Cells were in the stationary growth phase. Mean values and standard deviation of triplicates are shown.

### Influence of cell density and fluorogen concentration

We studied the influence of different cell concentrations and fluorogen concentrations on tdFAST2:HBR-3,5DOM fluorescence in *M. acetivorans*. With an increase in *M. acetivorans* cell concentration and a constant fluorogen concentration (5 µM), tdFAST2:HBR-3,5DOM fluorescence increased linear (Fig. 2a). Autofluorescence and OD(600) measured by the microplate reader increased similarly to tdFAST2:HBR-3,5DOM fluorescence (Fig. 2a) and, thus, could be used for normalization of tdFAST2:HBR-3,5DOM fluorescence values when making comparisons of different cultures.

**Figure 2.**
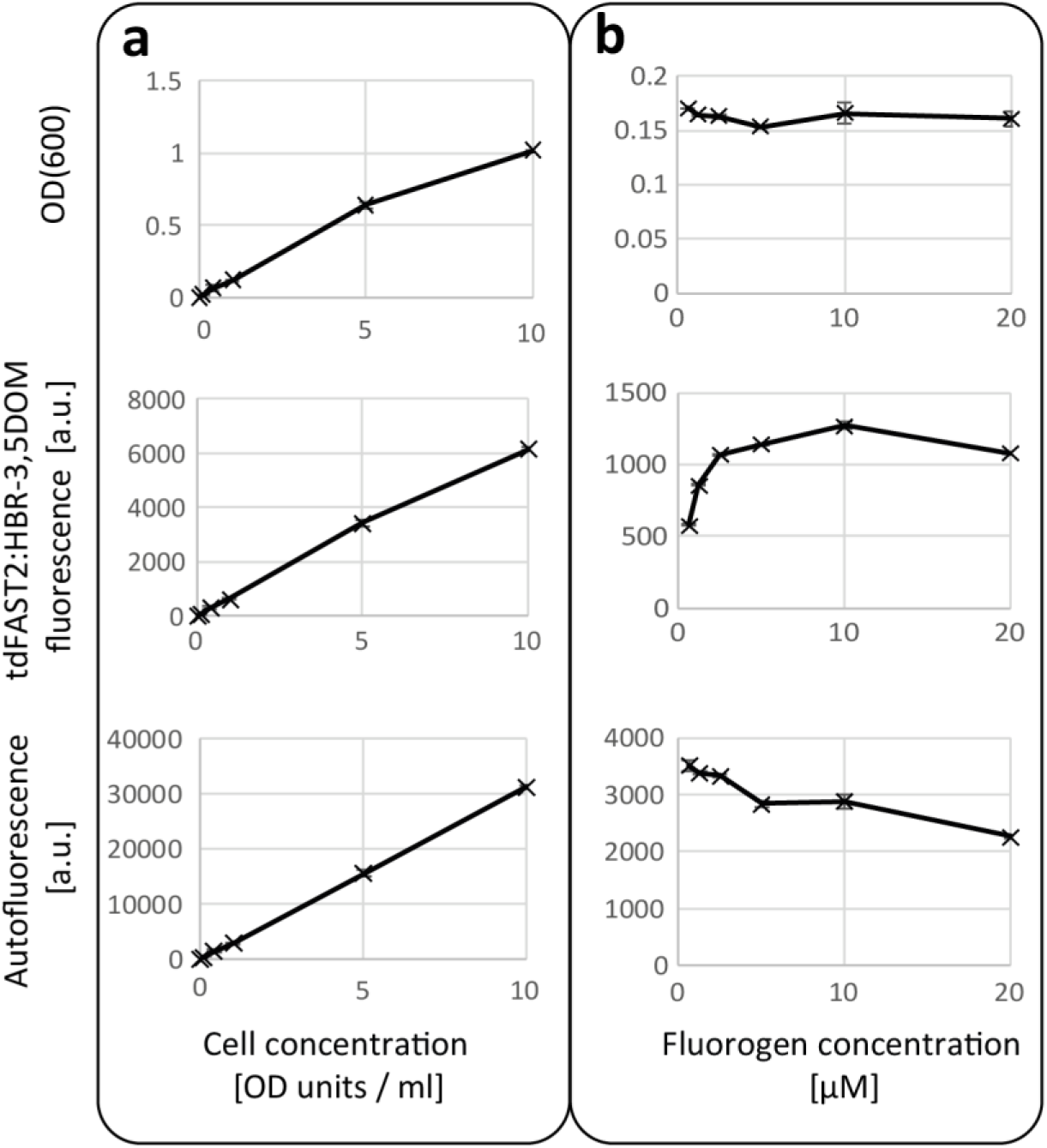
Wide ranges of cell concentration and fluorogen concentration can be used to measure tdFAST2:HBR-3,5DOM fluorescence in *M. acetivorans*. *M. acetivorans* cells expressing tdFAST2 were analysed in the exponential growth phase. A microplate reader was used to measure OD(600), tdFAST2:HBR-3,5DOM fluorescence (λ_Ex_ = 515 nm / λ_Em_ = 590 nm), and autofluorescence (λ_Ex_ = 420 nm / λ_Em_ = 480 nm). **a**) Correlation of fluorescence and cell-density when the fluorogen concentration (5 µM) is constant. **b**) Influence of the fluorogen concentration on the fluorescence when the cell concentration is constant (1 OD unit / ml). Note that, due to different light path lengths, the OD(600) determined by the microplate reader (upper panel) is smaller than the actual cell concentration (OD units / ml) which was determined using a standard spectrophotometer. Mean values and standard deviation of duplicates are shown.

In order to identify the optimal fluorogen concentration, the cell concentration was kept constant (1 OD unit / ml) while the fluorogen concentration increased from 0.625 µM to 20 µM. tdFAST2:HBR-3,5DOM fluorescence increased rapidly up to a concentration of 2.5 µM before reaching a plateau (Fig. 2b). Notably, higher fluorogen concentrations led to a decrease of autofluorescence but didn’t influence the OD(600) measured by the microplate reader. Based on these results, a fluorogen concentration of 5 µM was used for further experiments.

### tdFAST2:HBR-3,5DOM fluorescence during different growth phase

Following protein accumulation over time is an important application for fluorescent reporter systems. Therefore, we followed tdFAST2:HBR-3,5DOM fluorescence in *M. acetivorans* cells in different growth phases. tdFAST2:HBR-3,5DOM fluorescence was highest during late exponential growth phase while fluorescence was lowest in the stationary phase (Fig. 3). The *mcrB* promoter controlling tdFAST2 expression is considered a strong and constitutive promoter (18). Therefore, the differences in tdFAST2:HBR-3,5DOM fluorescence are more likely due to general, growth phase-dependent fluctuations in protein accumulation than due to promoter-specific differences. Consequently, when using tdFAST2:HBR-3,5DOM fluorescence for comparisons between cultures, they should be in the same growth phase. Remarkably, also autofluorescence was highest in the late exponential growth phase (Fig. 3) further supporting the hypothesis of growth phase-dependent differences of *M. acetivorans* cultures.

**Figure 3.**
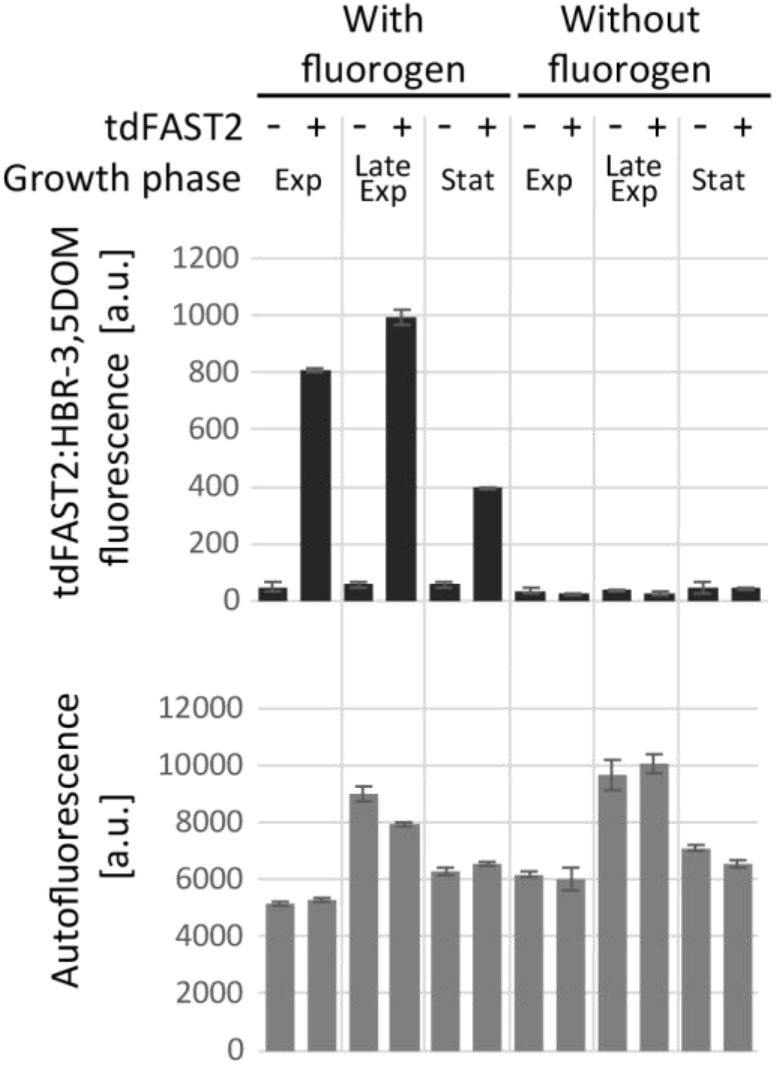
*M. acetivorans* fluorescence changes at different growth phases. *M. acetivorans* cells expressing tdFAST2 (+) or harbouring an empty vector construct (-) were analysed at exponential (Exp; OD(600) 0.3 – 0.6), late exponential (Late Exp; 0.8 – 1.1), and stationary growth (Stat; >1.1). tdFAST2:HBR-3,5DOM fluorescence (λ_Ex_ = 515 nm / λ_Em_ = 590 nm), and autofluorescence (λ_Ex_ = 420 nm / λ_Em_ = 480 nm) is shown. Mean values and standard deviation of duplicates are shown.

### Influence of tdFAST2 expression on multiplication

An important feature to consider when utilizing a reporter protein is its potential harmful/toxic effect on the cell. Therefore, we studied whether tdFAST2 expression has a general negative influence when expressed in methanogens. As shown in Fig 4, tdFAST2 expression leads only to a minor delay in cell growth and, thus, can be considered as non-toxic for methanogens.

**Figure 4.**
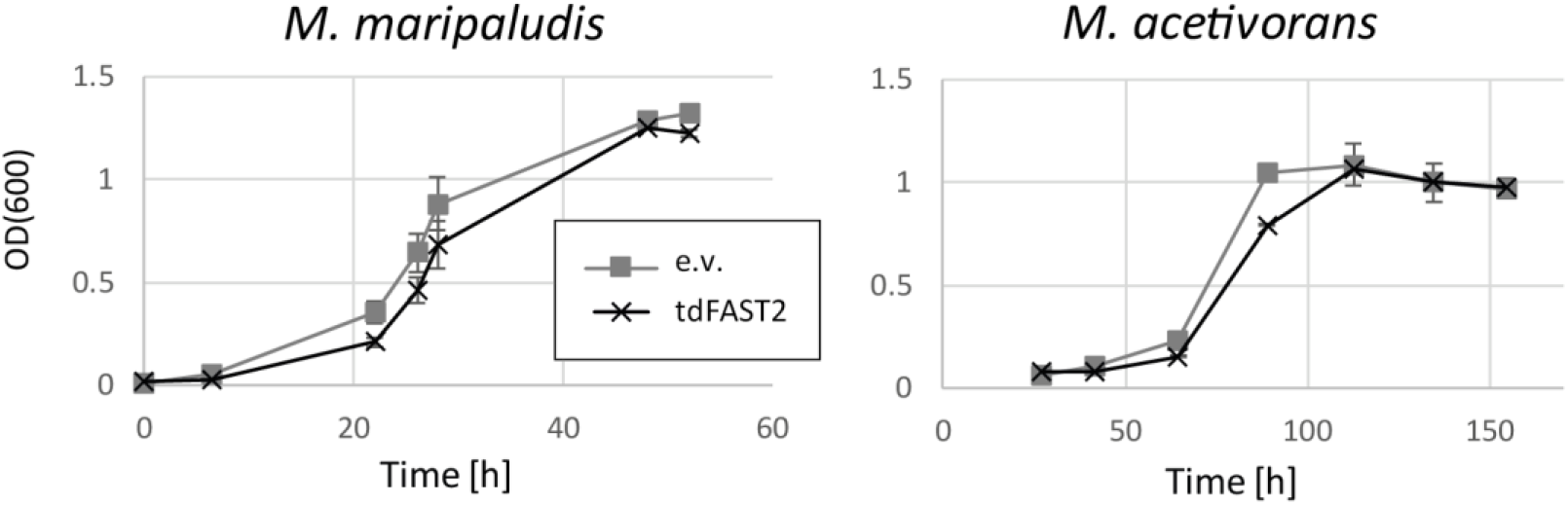
tdFAST2 is not toxic for methanogens. Growth of *M. acetivorans* and *M. maripaludis* expressing eighter tdFAST2 or harbouring an empty vector construct (e.v.) was analysed. Mean values and standard deviation of triplicates are shown.

### Flow cytometry

Detecting a reporter protein’s fluorescence via flow cytometry allows the detection of individual cells’ fluorescence and can be considered as gold standard for detection of a fluorescent reporter. We tested whether flow cytometry can be used to detect tdFAST2:HBR-3,5DOM fluorescence in methanogens. As shown in Fig. 5, tdFAST2:HBR-3,5DOM fluorescence was clearly detectable in tdFAST2-expressing *M. acetivorans* cells (Fig. 5a) and *M. maripaludis* cells (Fig. 5b) when HBR-3,5DOM was added. Fluorogen concentrations ranging from 2.5 to 10 µM were used and had a minor influence on the detected tdFAST2:HBR-3,5DOM fluorescence. This is in accordance with measuring the tdFAST2:HBR-3,5DOM fluorescence using a microplate reader (Fig. 2b). We also detected methanogens’ F420-dependent autofluorescence and found a little increase in fluorescence when cells showed a high tdFAST2:HBR-3,5DOM fluorescence. Probably, detection of autofluorescence detects a small amount of tdFAST2:HBR-3,5DOM fluorescence as well (Fig. 5a, b). When no tdFAST2 was expressed, fluorogen addition had no influence on the fluorescence detected.

**Figure 5.**
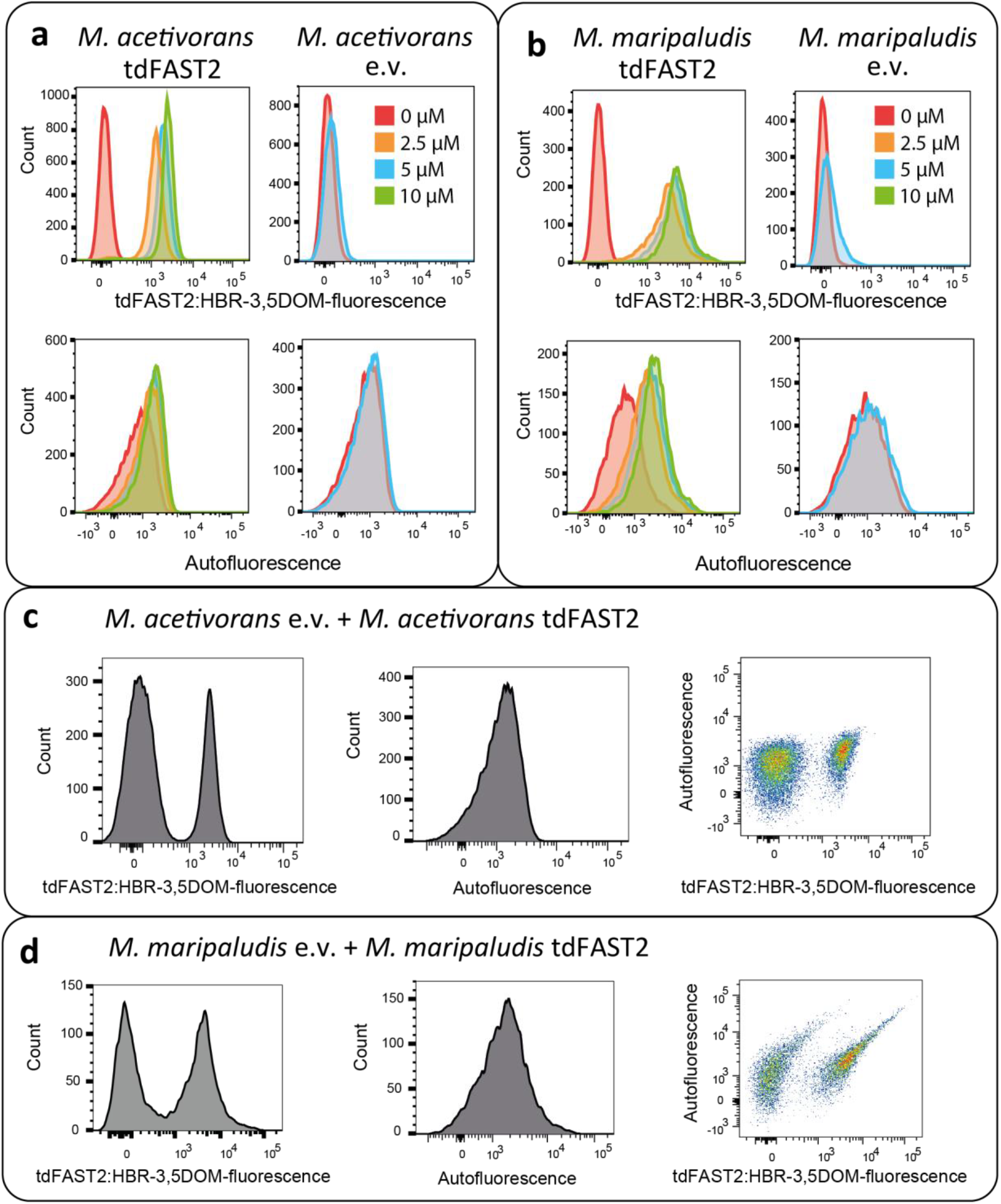
Flow cytometry allows visualization of tdFAST2 expression and separation of populations. Histograms of *M. acetivorans* (**a**) and *M. maripaludis* (**b**) cells expressing tdFAST2 (tdFAST2) or control cells carrying an empty vector construct (e.v.) in presence of increasing fluorogen (HBR-3,5DOM) concentrations. (**c** and **d**) Mixtures of tdFAST2-expressing cells and non tdFAST2-expressing cells were analyzed in the presence of 5 mM HBR-3,5DOM.

One application for flow cytometry is to identify distinct populations within mixed cultures. Thus, we measured *M. acetivorans* and *M. maripaludis* cultures in which tdFAST2-expressing cells were mixed with cells not harbouring the tdFAST2 gene. Flow cytometry with mixed cultures clearly identified two tdFAST2:HBR-3,5DOM fluorescence peaks (Fig. 5c and 5d). As expected, only one peak was detected for autofluorescence. Consequently, flow cytometry can be used to distinguish and separate tdFAST2-expressing cells from non-tdFAST2 expressing cells based on tdFAST2:HBR-3,5DOM fluorescence.

### Generation of Golden Gate cloning vectors for tdFAST2-tagged proteins

To simplify the use of tdFAST2 as fluorescence reporter in for *M. acetivorans*, we generated vectors that allow to generate N- or C-terminal tdFAST2 fusions in a one-step Golden Gate cloning reaction (19, 20). For this, the plasmid pNB730 was domesticated by removing *Bsa*I sites in the vector backbone and the tdFAST2-encoding sequence was inserted. Additionally, a *lacZ* cassette was inserted to allow blue-white selection of clones. The *lacZ* cassette is flanked by *Bsa*I restriction sites and can be replaced by the protein sequence of interest. For cloning, the sequence of interest can be amplified with primers adding flanking BsaI sites and the corresponding overhangs (AATG and AAGC) for insertion (forward primer overhang: 5’-TTTGGTCTCTAATG; reverse primer overhang: 5’-TTTGGTCTCTAAGC). If the inserted sequence is in the same open reading frame as tdFAST2, a AGGGSGGG (C-terminal tdFAST2) or GGGSGGGM (N-terminal tdFAST2) linker will be encoded between tdFAST2 and the protein of interest (Fig. 6). The generated plasmids pMaFAST(C) and pMaFAST(N) contain a ΦC31 attB sites allowing integration into the *M. acetivorans* genome at attP sites as well as the *pac* and *bla* genes for selection in *M. acetivorans* and *E. coli*, respectively.

**Figure 6.**
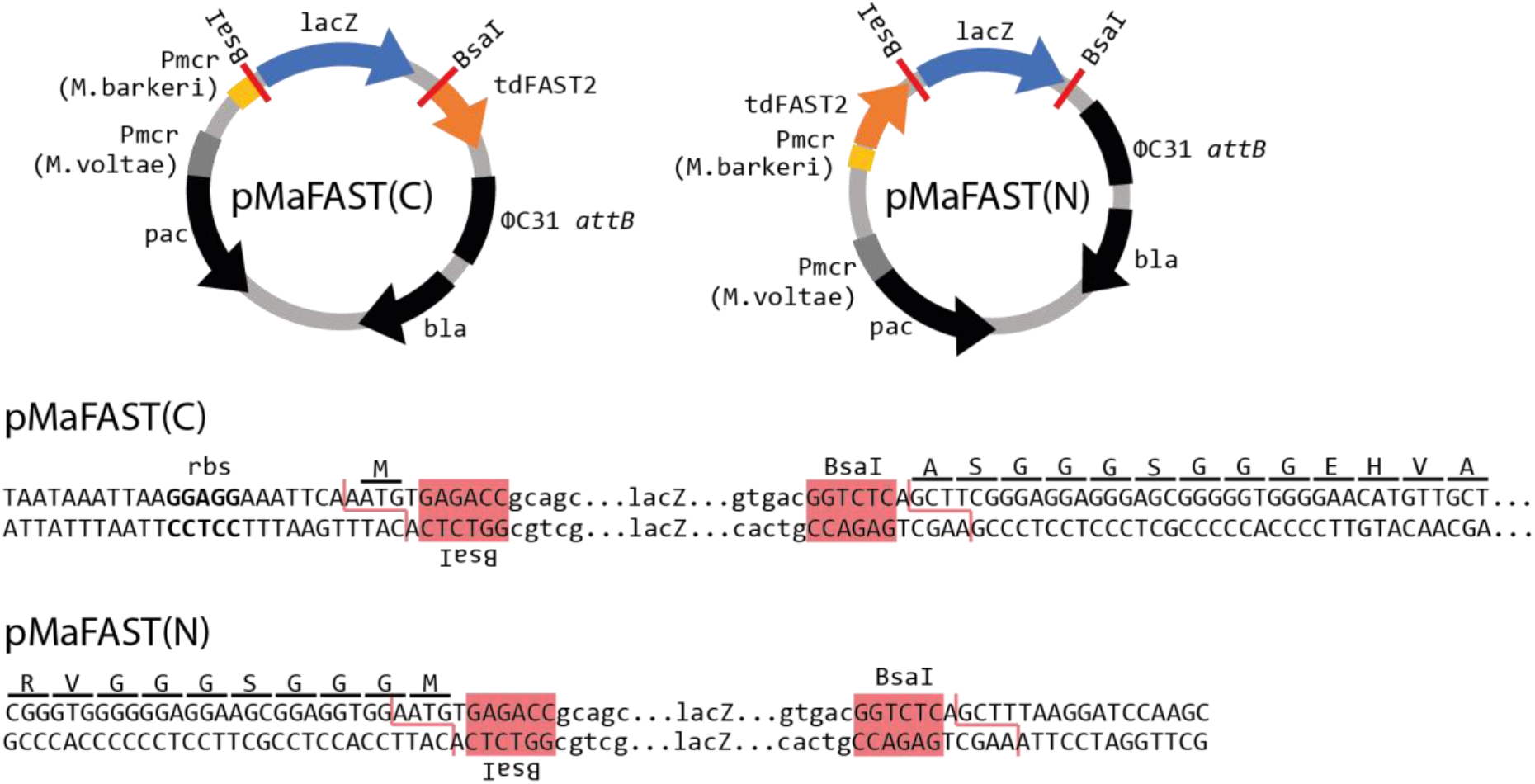
Golden Gate cloning vectors to study tdFAST2-tagged proteins. The plasmids pMaFAST(C) and pMaFAST(N) encode the tdFAST2 gene which is codon-optimized for expression in *Methanosarcina*. Golden Gate cloning (*Bsa*I) can be used to replace the *lacZ* cassette with a protein-encoding sequence of interest leading to a C- or N-terminal fusion to the tdFAST2. Nucleotide sequence of the cloning sites are shown below. The *Bsa*I sites, ribosome-binding site (rbs) and the tdFAST2 open-reading frame is given.

## Discussion

Here, we introduce tdFAST2:HBR-3,5DOM as reliable fluorescent reporter system for the two methanogenic model organisms *M. acetivorans* and *M. maripaludis*. tdFAST2:HBR-3,5DOM fluorescence can be measured with a microplate reader as well as by flow cytometry and shows no overlap with methanogens’ autofluorescence, which can be measured in parallel. Importantly, tdFAST2 expression has no cytotoxic effect. A washing step was introduced to remove background fluorescence from the medium. This goes along with recent findings reporting a strong HBR-3,5DOM-dependent background fluorescence in *M. maripaludis* cultures (14). Similarly, a washing step also reduced background fluorescence from culture medium when FAST:fluorogen fluorescence was studied in anaerobic *Clostridium* (21).

### A new reporter system for methanogens

Previously, the most frequently used reporter in methanogenic archaea was *uidA. uidA* encodes β-glucuronidase which can be quantified by *in vitro* activity assays using cell extracts from *Methanosarcina* spp. (22, 23) as well as *Methanococcus* spp. (24). Less commonly used enzymatic reporters in methanogens include acetohydroxyacid synthase, β-Galactosidase, and β-Lactamase (25, 26). Disadvantages of enzymatic reporters include i) the need for multiple handling steps hindering high throughput screenings, ii) the inability to study single cells, and iii) the need of relatively high cell culture volumes. These disadvantages were partly overcome by using the fluorescent reporter mCherry in *M. maripaludis* (27, 28). Although mCherry can be easily monitored using microplate readers and shows no overlap with methanogens’ autofluorescence, its use is time-consuming and includes cell lysis by freeze-thawing and overnight exposure to oxygen to allow maturation of the chromatophore. The latest development of a new reporter system for methanogens is the quantification of protein accumulation in *M. maripaludis* using FAST1:HMBR fluorescence (14). There, FAST1 was N-terminally fused to FruA, a hydrogenase subunit, which is known to be more abundant upon growth in formate-containing medium compared to hydrogen-containing medium. FAST1-FruA abundance in formate-grown and hydrogen-grown cells was quantified by anoxic microscopy in which FAST1:HMBR fluorescence of hundreds of single cells was monitored. Quantification of fluorescence intensities of several hundred single cells via microscopy is time consuming and impractical and, unfortunately, it remains unclear why FAST1:HMBR fluorescence in FAST1-FruA expressing cultures was not quantified using a microplate reader. Based on our experiences, the use of tdFAST2:HBR-3,5DOM fluorescence in *M. acetivorans* and *M. maripaludis* outcompetes the above-mentioned reporter systems to study promoter activity/protein accumulation and, by using flow cytometry, even allows the accurate and efficient quantification of individual cells’ fluorescence.

### Aerobic vs. anaerobic detection of FAST:fluorogen fluorescence

Due to technical reasons, we performed fluorescence measurements solely under aerobic conditions. Notably, also most enzymatic reporter systems are performed aerobically and the use of mCherry in *M. maripaludis* even includes over-night exposure to oxygen prior to protein quantification (see above). For quantification of tdFAST2:HBR-3,5DOM fluorescence, cells were harvested from anaerobic cultures, pelleted, washed, and subsequently analyzed using a plate reader or by flow cytometry. Handling time before fluorescence measurements was short (5 – 10 min) and even longer waiting times before addition of 3,5DOM (up to 1 h) had no influence on the fluorescence (data not shown). We conclude that it is not necessary to use anaerobic conditions when studying tdFAST2:fluorogen fluorescence in anaerobic methanogens. The procedure of analyzing FAST:fluorogen fluorescence in anaerobic organisms under aerobic conditions was used before. For example, FAST:HMBR fluorescence of anaerobic *Clostridium* organisms was monitored aerobically using confocal microscopy, microplate reader measurements and flow cytometry measurements (21).Another recent study applied flow cytometry (aerobic) to detect fluorescence of anaerobic, acetogenic bacteria (29). Other studies avoid oxygen when detecting FAST:fluorogen fluorescence of anaerobic organisms, e.g., by placing a fluorescence microscope or microplate reader into anaerobic chambers (14, 30). To our knowledge, no severe differences between detection of FAST:fluorogen fluorescence of anaerobic organisms under anaerobic and aerobic conditions were reported. We hypothesize that, although oxygen has a severe negative influence on the physiology of strictly anaerobic organisms, its short-term influence on protein abundance, protein localization and protein-protein interaction remains marginal. This assumption, however, awaits to be systematically analyzed in future studies.

### Applications of tdFAST2:HBR-3,5DOM fluorescence in methanogens

tdFAST2:HBR-3,5DOM fluorescence allows multiple applications including *in vivo* protein localization and the analysis of *in vivo* protein-protein interaction when the FAST2 protein is split and fused to different proteins of interest. Both applications were recently successfully performed in *M. maripaludis* using a FAST-fluorogen fluorescence (14).

Flow cytometry allows high-throughput measurement of multiple physical and biochemical characteristics of cells (31). The ability to detect and sort a large quantity of individual cells based on tdFAST2:HBR-3,5DOM fluorescence opens new routes to study and engineer methanogens. For example, differential fluorescence induction (DFI) strategies (32) can be used to identify inducible promoters for *Methanosarcina* and *Methanococcus* species. For this, a complex promoter library could be screened by cloning the library in front of a promoterless tdFAST2 gene and transformation into methanogens. Subsequently, the transformed culture could be treated with the desired inducer for gene expression and individual cells would be screened and sorted for high tdFAST2:HBR-3,5DOM fluorescence using fluorescence-activated cell sorting (FACS). In order to eliminate constitutive promoters, cells with a high tdFAST2:HBR-3,5DOM fluorescence could be re-grown and sorted again without the inducer. Obviously, this approach requires to perform cell sorting under anaerobic conditions to keep methanogens viable; a challenge that was successfully tackled before (33–36). DFI was successfully used in various microbes including *Salmonella, Streptococcus, Pseudomonas* and *Bacillus* species (32).

Another application of flow cytometry experiments using tdFAST2:HBR-3,5DOM fluorescence in methanogens is the ability to study population dynamics in mixed methanogen cultures to variations within a culture. Currently, the typical molecular characterization of *M. acetivorans* and *M. maripaludis* includes genetically modification (deletion or insertion) and comparing the modified cells to the wildtype. Usually, wildtype cells and modified cells are separated and characterized individually. Expression of tdFAST2 in either wildtype or modified cells allows studying mixed cultures where tdFAST2:HBR-3,5DOM fluorescence is used to separate both population. This real-time investigation of dynamics within mixed cultures allows conclusions about the modification’s relevance for the organism. Measuring population dynamics in mixed culture also gains more relevance in biotechnology where mixed cultures are increasingly used for production (37). Various methanogens are known to be involved in direct interspecies electron transfer (DIET) in co-culture with electron-donating bacteria (38, 39). The ability to separate two methanogen strains by using tdFAST2:HBR-3,5DOM fluorescence might become a relevant tool to study DIET co-cultures.

## Material and methods

### Strains, cultivation and transformation

The strains *M. acetivorans* WWM73 (18) and *M. maripaludis* S0001 (40) were used. Methanogens were cultivated under strictly anaerobic conditions. For *M. acetivorans* growth, high salt (HS) medium (41) was used containing 125 mM methanol (MeOH) or 50 mM trimethylamine (TMA) as the growth substrate. *M. maripaludis* was cultivated using McFC medium (42). PEG-mediated transformation (43, 44) was used for plasmid integration into methanogens and 2 µg/ml puromycin was used as selection marker when required. Plasmids pNB730::tdFAST2 and pNB730 were transformed in *M. acetivorans* and plasmids pMEV4::tdFAST2 and pMEV4 were transformed in *M. maripaludis*.

For plasmid construction, *Escherichia coli* strain TOP10 (Thermo Fisher Scientific) was used employing standard methods and enzymes purchased from Thermo Fisher Scientific or New England BioLabs (NEB).

### Cloning of tdFAST2

The tdFAST2 (also known as td-iFAST (45)) sequence was codon optimized using the Eurofins GENEius (www.geneius.de) codon-optimization tool and the *M. maripaludis* S2 and *M. acetivorans* C2A codon usage tables taken from http://www.kazusa.or.jp/codon/.A GGGSGGG linker connecting the two FAST2 domains was used and internal restriction sites (*Bsa*I, *Bbs*I and *Msm*BI) were not allowed. For expression in *M. maripaludis*, a ribosome-binding site (AGTGGGAGGTGCGC) and the transcriptional terminator from MMP1100 (AAATTCTTCTTCTTTTAAACGTTCTCCAGT (46)) was attached to the tdFAST2-coding sequence. For cloning, sequences were flanked by *Nde*I and *Bam*HI sites or *Spe*I and *Pst*I, respectively. Sequences were synthesized from Eurofins Scientific and are given in Supplementary file 1. Classical cloning was used to clone the synthesized fragments into pBN730 (47) for expression in *M. acetivorans* and pMEV4 (48) for expression in *M. maripaludis*. The generated plasmids were named pNB730::tdFAST2 and pMEV4::tdFAST2.

### Fluorescence measurements

For microplate reader measurements, a BioTek Cytation 3 Microplate Reader was used. Cells were harvested from anaerobic cultures and the optical density at 600 nm (OD600) was determined using a spectrophotometer. Cells were pelleted by centrifugation (11000 g, 2 min) and resuspended in a salt solution (400 mM NaCl, 13 mM KCl, 54 mM MgCl_2_, 2 mM CaCl_2_) mimicking the HS medium. The high salt content serves to prevent cell lysis due to osmotic changes. Cells were pelleted again and subsequently resuspended in salt solution containing the fluorogen. If not stated otherwise, a fluorogen concentration of 5 µM and a volume to obtain a cell density of 1 OD(600) unit/ml was used. 100 µl aliquots were analysed in Nunc 96-well flat-bottom microplates. Salt solution was used to determine background fluorescence values which was subtracted from the cells’ fluorescence.

For flow cytometry, a LSRFortessa Cell Analyzer (BD Biosciences) was used. Cells were harvested from anaerobic cultures, pelleted by centrifugation (11000 g, 2 min), and resuspended in salt solution. The fluorogen HBR-3,5DOM (45) was added immediately before flow cytometry analysis. For tdFAST2:HBR-3,5DOM fluorescence, a blue laser (488 nm excitation) and 610/20 nm filter were used. For autofluorescence, a violet laser (405 nm excitation) and 510/50 nm filter were used, and 20,000 events were recorded. The flow cytometry analysis was performed at the HiLife Flow Cytometry Unit, University of Helsinki. Results were analyzed using the FlowJo™ v10.8 Software (BD Biosciences). 4-hydroxy-3,5-dimethoxybenzylidene rhodanine (HBR-3,5DOM) and 4-hydroxy-3-methylbenzylidene rhodanine (HMBR) were purchased from Twinkle Bioscience (www.the-twinkle-factory.com) and stored as a 5 mM stock solution in DMSO at -20°C.

### Generation of golden gate cloning vectors

The vector pNB730 was turned into the Golden Gate cloning destination vectors pMaFAST(C) and pMaFAST(N) using the protocol “Accommodating a vector to Golden Gate cloning” (19). For this, the *lacZ* cassette was amplified from pUC19 (New England Biolabs, cat. no. N3041S) using the primers oNA311 (ttgaagacaaAATGtgagaccgcagctggcacgacaggtttc) and oNA312 (ttgaagacaaAAGCtgagaccgtcacagcttgtctgtaagcg). The primer overhangs add *Bpi*I restriction sites (gaagac), 4-nt fusion sites and *Bsa*I sites (ggtctc) in reverse complementary orientation to the *lacZ* fragment. For generation of pMaFAST(C) the vector backbone was amplified in two parts from pNB730::tdFAST2 to remove one internal *Bsa*I site. For this, primer pairs oNA323 (ttgaagacaaGCTTTAAgGATCCAAGCTTGGGCCCTCG) / oNA304 (ttgaagacaaACGCTCACCGGCTCCAGATTTATC) and oNA305 (ttgaagacaaGCGTGGATCTCGCGGTATCATTG) / oNA328 (ttgaagacaaCATTCCACCtCCGCTtCCTCCcCCCACCCGTTTTACAAACACCCAGTAAC) were used. For generation of pMaFAST(N), the vector backbone was also amplified in two parts from pNB730::tdFAST2 using the primer pairs oNA343 (ttgaagacaaGCTTCGGGAGGGAGCGGGGGTGGGGAACACGTCGCGTTTGGCTC) / oNA304 and oNA305 / oNA318 (ttgaagacaaCATTTGAATTTCCTCCTTAATTTATTAAAATCATTTTGGGAC) were used. Primers have *Bpi*I restriction sites with specific 4-nt fusion sites to be compatible to each other upon ligation. Two backbone fragments and the *lacZ* fragment were fused together in a Golden Gate cloning reaction with *Bpi*I. The cloning product was transformed into *E. coli* and plated on ampicillin- and X-gal-containing LB plates. Blue colonies were selected and verified by sequencing the areas around the *Bsa*I cloning sites.

## Acknowledgement

We thank Prof. N. Buan for providing the plasmid pNB730 and Prof. W. Whitman for providing pMEV4. This work was supported by grants from Novo Nordisk Foundation and the Academy of Finland (grant NNF19OC0055464 to N.A. and grants NNF19OC0054329 and 326020 to S.S.).

## Figure legends

**Supplementary Figure 1.**
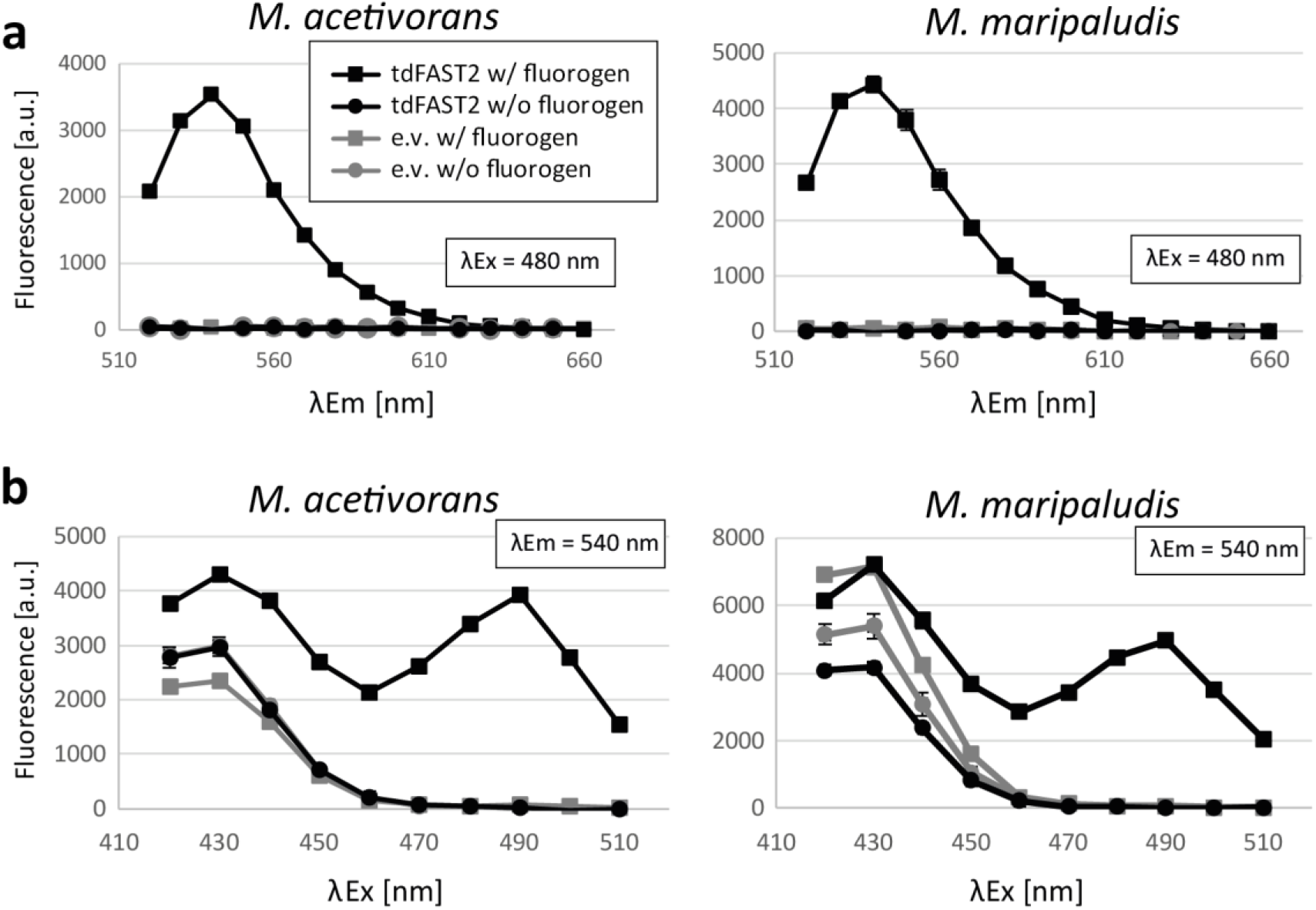
tdFAST2-expressing methanogens show a specific fluorescent when the fluorogen HMBR is present. Fluorescence of *M. acetivorans* and *M. maripaludis* cells is shown. **a**) Fluorescence spectrum upon excitation at 480 nm. **b**) Fluorescence at 540 nm when different excitation wavelengths are applied. Cells expressing tdFAST2 (tdFAST2) or control cells carrying an empty vector construct (e.v.) were analysed in presence (w/ fluorogen) or absence (w/o fluorogen) of HMBR. Used cells were in the stationary growth phase. Mean values and standard deviation of duplicates are shown.

## Appendix

**Supplementary File 1.**
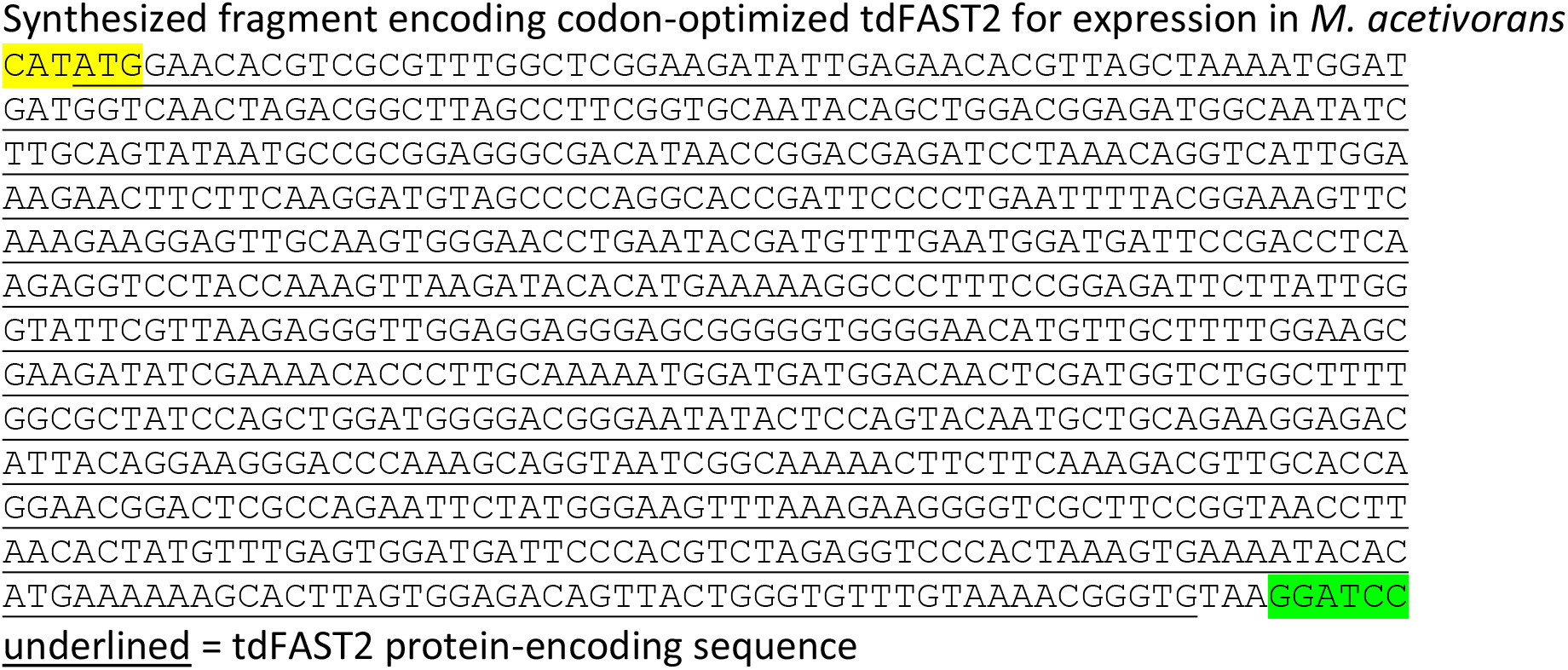
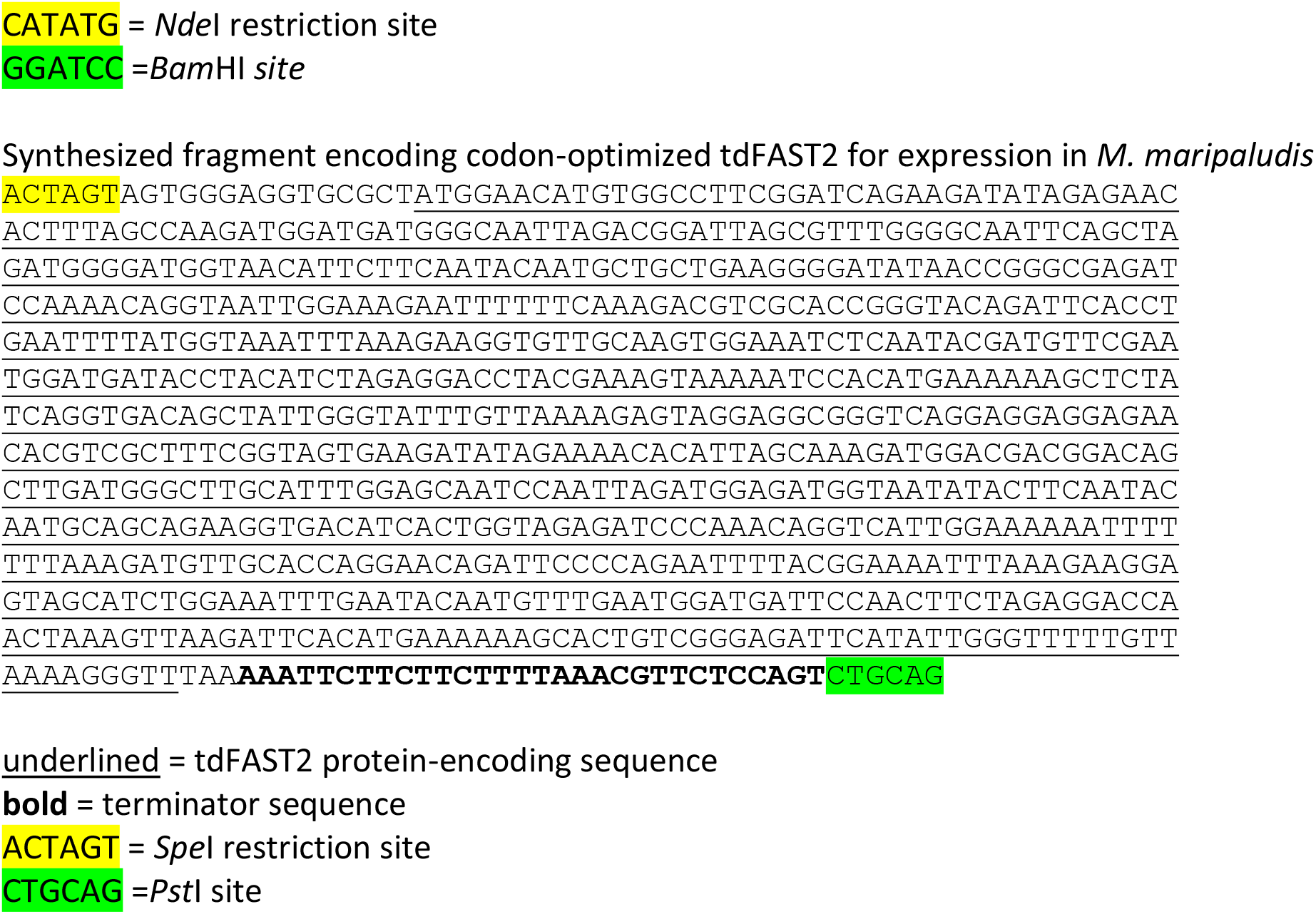
Synthesized sequences.

**Supplementary File 2.**
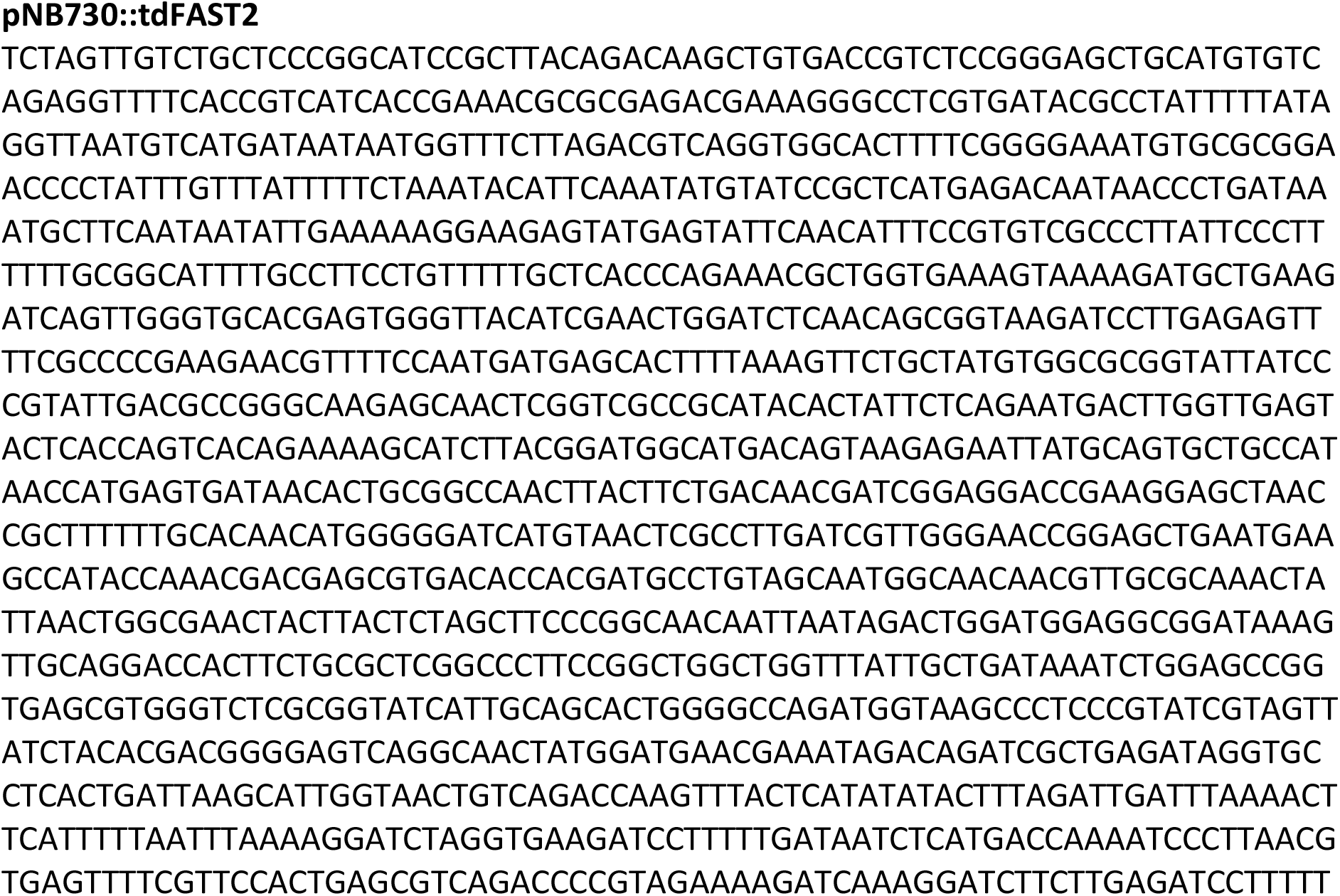
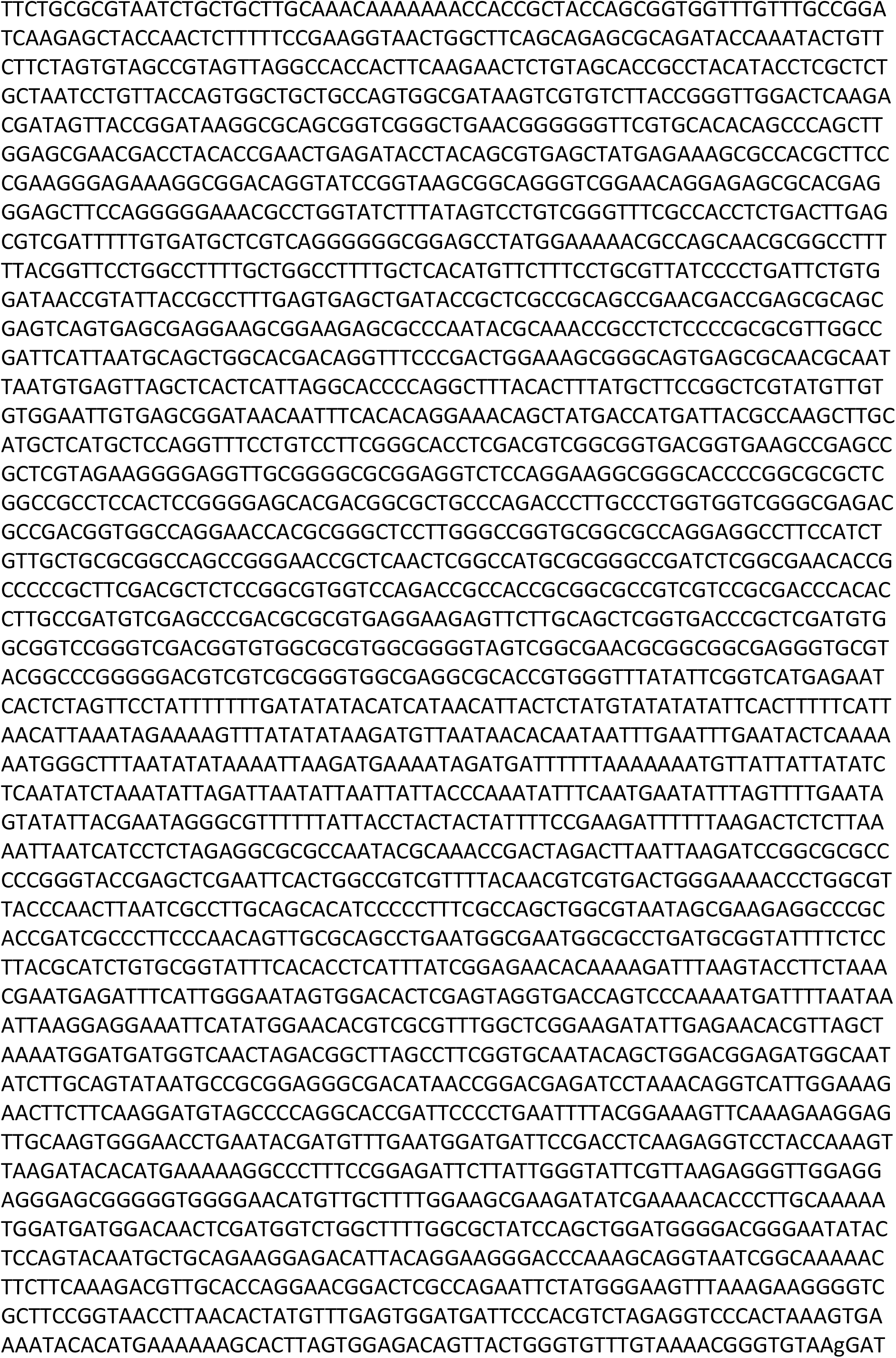
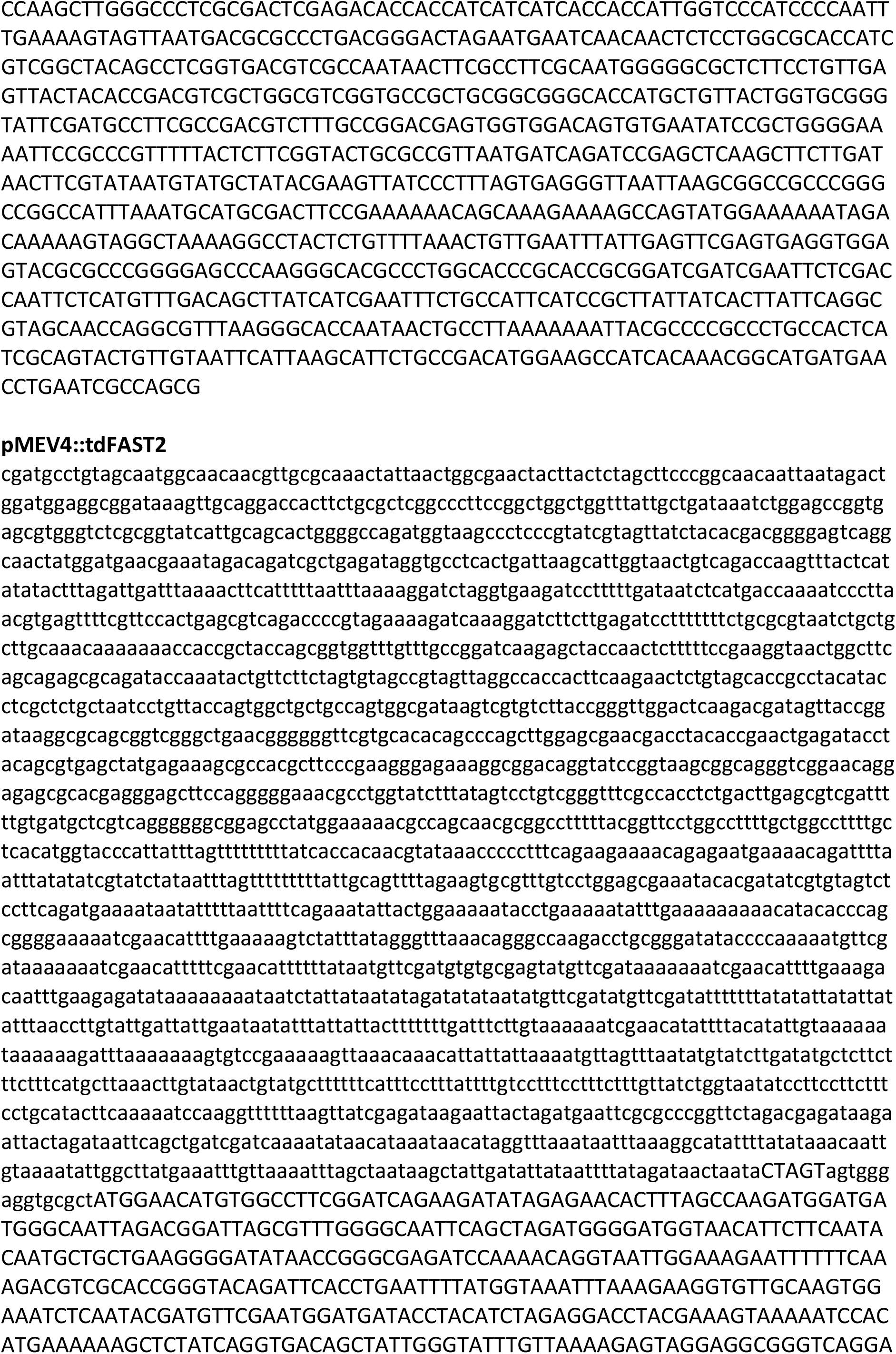
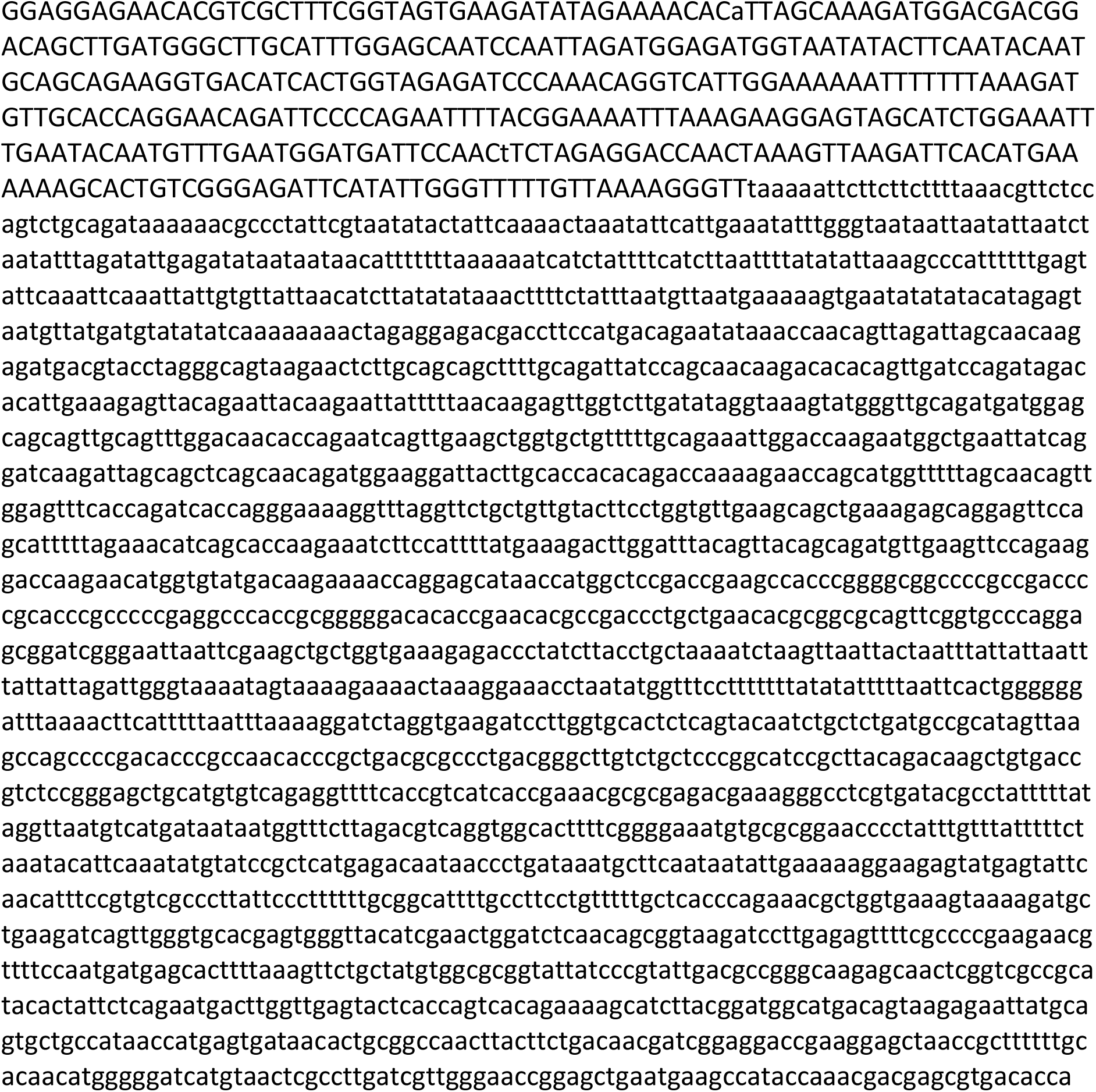
Plasmid sequences.

